# From Black Box to Biological Insight: AttentioFuse Unlocks Multi-Omics Dynamics in Lung Cancer

**DOI:** 10.1101/2025.07.23.665084

**Authors:** Yuhang Huang, Yungang He, Lei Liu, Fan Zhong

**Affiliations:** Institutes of Biomedical Sciences, Fudan University, 131 Dongan Road, Shanghai, Shanghai 200032, China; Intelligent Medicine Institute, Fudan University, Shanghai 200032, China; Shanghai Institute of Infectious Disease and Biosecurity, Fudan University, Shanghai 200032, China; Shanghai Institute of Stem Cell Research and Clinical Translation, Shanghai 200120, China

**Keywords:** interpretable deep learning framework, mid-fusion strategy, multi-omics, hierarchical attention mechanisms, NSCLC

## Abstract

Lung adenocarcinoma (LUAD) and squamous cell carcinoma (LUSC), the major subtypes of non-small cell lung cancer (NSCLC), exhibit distinct molecular landscapes that demand precision in prognosis and therapy. While deep learning models achieve high predictive accuracy, their “black-box” nature limits clinical translation. To address this, we propose AttentioFuse, an interpretable deep learning framework employing a Reactome-guided mid-fusion strategy for multi-omics integration. AttentioFuse innovates through three pillars: (1) dual-phase learning to preserve omics-specific patterns via independent sub-networks, (2) hierarchical attention mechanisms (cross-omics, feature-level, and fusion-layer) to dynamically quantify layer contributions, and (3) integrated explainability combining DeepSHAP and global attention weights for gene-to-pathway interpretation. Evaluated on TCGA LUAD/LUSC cohorts, AttentioFuse matches state-of-the-art performance in TNM staging, while uncovering actionable biological insights. The framework validates pan-NSCLC mechanisms like AKT/mTOR metabolic control and histology-divergent Notch signaling roles, while revealing novel pathways—developmental reactivation (T-stage), microbiota-driven metastasis (M-stage), and ECM remodeling—providing testable hypotheses for progression and personalized therapy. Crucially, AttentioFuse bridges computational predictions to clinical practice by proposing molecularly-guided combination therapies. This paradigm shifts toward interpretable-aware AI advances oncology by transforming black-box predictions into biologically grounded decision-support tools.

## Introduction

Lung adenocarcinoma (LUAD) and squamous cell carcinoma (LUSC) collectively represent over 80% of non-small cell lung cancer (NSCLC) cases, yet their distinct molecular landscapes pose persistent challenges for precision prognosis [1]. Current deep learning approaches, while achieving respectable predictive accuracy in survival analysis, often function as “black boxes” that obscure the biological rationale behind their predictions—a critical limitation for clinical translation [2].

Traditionally, deep neural networks have delivered high predictive accuracy, but they are often criticized for their lack of interpretability. Recent efforts employing early data fusion strategies (IntegratalNet) have shown that integrating multiple data modalities can significantly enhance survival prediction in NSCLC. For instance, Ellen et al. [3] developed a multimodal integration approach that uses denoising autoencoders to compress and fuse diverse data types (e.g., mRNA, miRNA, and DNA methylation) at an early stage, demonstrating that models combining multiple modalities outperform those based on a single one. Similarly, Elbashir et al. [4] adopted a comparable strategy by replacing the elastic net with a graph attention network (GAT) [5], resulting in superior survival prediction metrics. One notable model in this domain is P-NET [6], an IntegratalNet that achieves strong interpretability by incorporating prior biological knowledge via mask matrices. This design reveals key molecular and pathway features, as validated through experimental studies.

However, such methods may overlook the distinct biological characteristics of each omics layer and fail to provide insight into the relative contributions of genomic alterations to disease progression. This study refines P-NET’s core ideas by shifting from a potentially early fusion approach to a mid-stage fusion strategy combined with an attention fusion layer [7]. We propose an interpretability-oriented mid-fusion framework AttentioFuse (Figure 1) with three key innovations:

**Figure 1.**
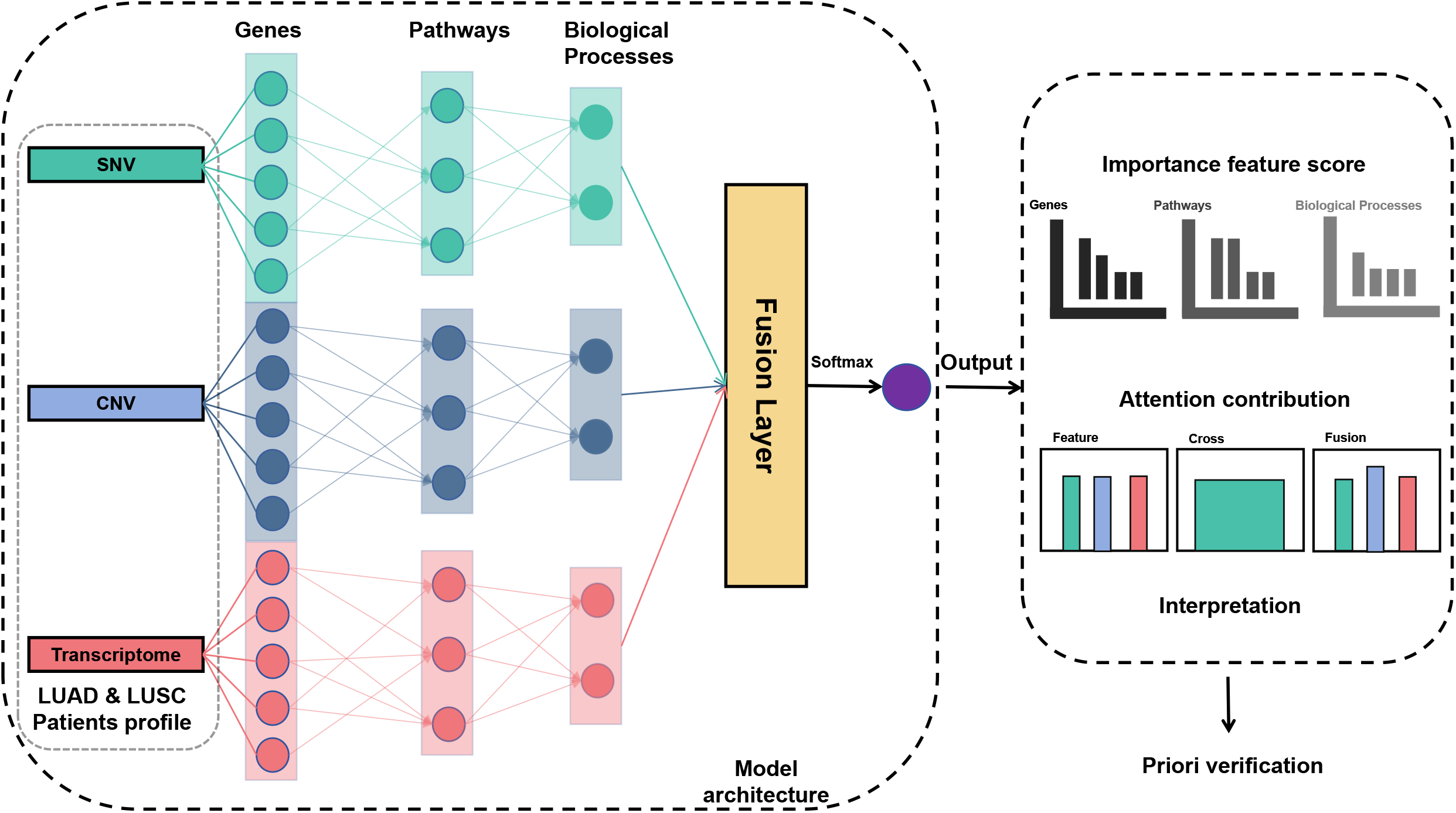
Architecture of the AttentioFuse Framework. The AttentioFuse framework processes multi-omics data (SNV, CNV, Transcriptome) through individual sub-networks to preserve omics-specific patterns. These layer-specific representations are then fused at a mid-stage using a trainable attention layer, weighting each omics layer’s contribution. The model integrates interpretation by combining DeepSHAP with global attention weights, providing both feature importance scores at the gene and omics levels and attention contribution visualizations for interpretation and subsequent experimental validation.

Dual-phase learning: Preserves omics-specific patterns through independent sub-net branches prior to cross-modal fusion.

Dynamic attention weighting: Quantifies layer-specific contributions via trainable attention gates.

Integrated explainability: Combines DeepSHAP’s local interpretation with global attention weights, reconciling feature importance at both gene and omics levels [8].

Our approach prioritizes biological insight over marginal performance gains. Benchmark tests on The Cancer Genome Atlas (TCGA) data [9] show comparable accuracy to state-of-the-art models, and with significantly enhanced interpretability. This paradigm shifts from “performance-centric” to “interpretation-aware” modeling provides clinicians with actionable biological hypotheses, bridging the critical gap between algorithmic predictions and therapeutic decision-making.

## Methods

### Data Acquisition and Processing

This study leveraged publicly available data from TCGA for LUAD and carcinoma LUSC. Data acquisition utilized the TCGAbiolinks package (downloaded September 2024) [10]. Transcriptomic profiling, copy number variation (CNV), and single nucleotide variation (SNV) datasets were obtained for both LUAD and LUSC cohorts.

For transcriptomic profiles, raw TPM-normalized expression matrices underwent mitochondrial gene removal to reduce potential noise from highly variable mitochondrial transcripts, yielding processed datasets of 600 samples in LUAD and 562 samples in LUSC. CNV data derived from GISTIC v2.0 analysis [11] were filtered to retain high-confidence alterations (high-level amplifications/deep deletions), producing final samples of 518 in LUAD and 504 in LUSC. SNV processing involved three-tier filtering: (1) exclusion of non-functional variants (silent/UTR/intron/IGR), (2) quality control with alternative allele depth > 5, and (3) deduplication by sample-gene pairs, resulting in 556 samples in LUAD and 544 samples in LUSC mutation matrices.

Pathological staging labels were curated from TCGA clinical annotations through a standardized binarization protocol. Primary tumor (T), regional lymph nodes (N), and distant metastasis (M) stages [12] were classified into clinically informative low (L) and high (H) risk categories. For tumor (T) stage, low risk encompassed Tis and T1-2, while high risk was T3-4. For node (N) stage, low risk was N0, and high risk was N1-3. For metastasis (M) stage, low risk was M0, and high risk was M1.

### Missing Value Imputation

To address structural missingness while preserving biological relevance, we implemented a density-based hybrid imputation strategy. The process commenced with feature-wise mean initialization to establish baseline values, followed by DBSCAN [13] clustering (*ε*=0.5, *min_samples*=5) on standardized data (*μ*=0, *σ*=1) to identify local sample neighborhoods. Within each cluster, missing values were imputed using intra-cluster feature means, reverting to global means when cluster-specific data was unavailable. Samples failing to cluster (deemed noise) received global mean imputation. This tiered approach maintained global data distributions, evidenced by Kullback-Leibler divergence <0.05 relative to raw data [14], while leveraging local structure to enhance biological plausibility.

### Class-Balanced Augmentation

To mitigate TNM stage imbalance, borderline-SMOTE [15] synthesized minority class samples through borderline instance identification and *k*-nearest neighbor (*k*=5) interpolation. The algorithm selectively targeted minority samples near decision boundaries, generating synthetic instances via linear combinations of high-risk neighbors. Augmentation intensity was dynamically calibrated per TNM category, constrained to <15% synthetic samples in final training sets through probabilistic sampling. This threshold prevented overfitting while ensuring adequate representation of rare stages.

### Model design

AttentioFuse employing mask matrices derived from the Reactome database (v86) [16]. These mask matrices dictate the network’s layer configuration, with the number of nodes in each layer (Figure 2a), corresponding to the number of genes/features associated with each level of the Reactome hierarchy after data cleaning and feature selection.

**Figure 2.**
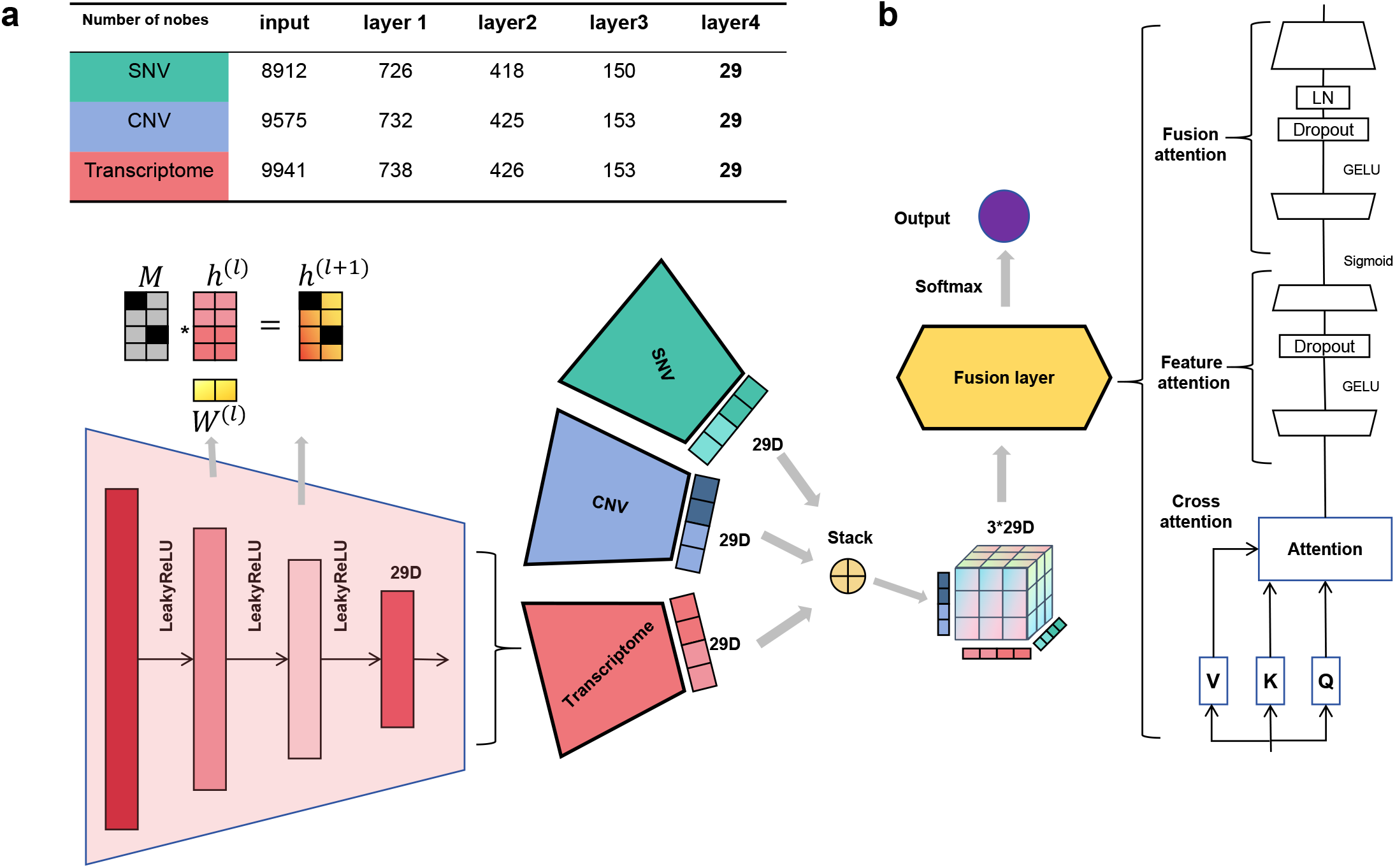
AttentioFuse Model Architecture and Interpretation Methodology. (a) Table showing the number of nodes in each layer of the sub-networks for each omics data type. (b) Detailed view of the mid-fusion stage, highlighting the stacking of omics representations, the fusion layer with attention mechanisms, and the interpretation framework. The “Feature attention” block refers to the attention mechanism used to calculate modality-specific pathway importance scores, while “Fusion attention” and “Cross attention” depict the components involved in cross-omics integration and pathway impact score derivation, combining attention weights and gradient attributions for model interpretation. Left panel (bottom) illustration of the LeakyReLU activated dense layers within each omics-specific sub-network.

**Figure 3.**
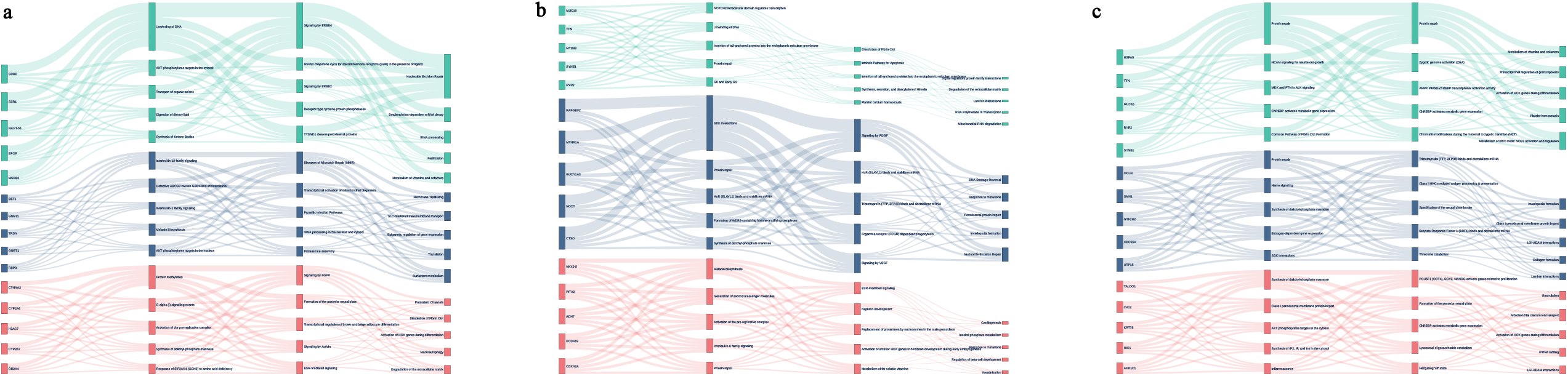
Sankey Diagram Visualization of Key Pathway Contributions to TNM Staging in LUSC. Sankey diagrams illustrate the pathway contributions to (a) M, (b) N, and (c) T-stages, respectively. Nodes are colored by omics data type: Transcriptome (red), CNV (blue), and SNV (green). The diagram highlights validated mechanisms such as Hippo-Notch signaling (T-stage), PDGFR/VEGF-metalloregulation (N-stage), and ERBB-HSP90 coupling (M-stage), alongside novel findings including developmental pathway reactivation and microbiota-driven metastasis, providing a visual summary of pathway importance across TNM stages in LUSC.

**Figure 4.**
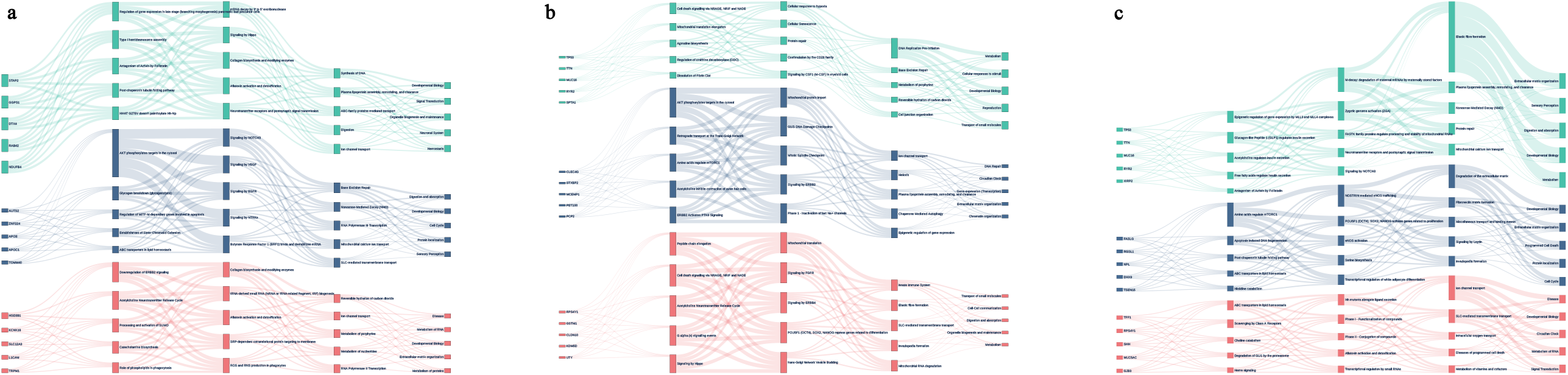
Sankey Diagram of Validated and Novel Pathway Contributions in LUAD. The diagram highlights validated mechanisms such as mTORC1-mediated metabolic control in T-stage and AKT-VEGF angiogenic coupling in M-stage, alongside novel insights including embryonic mRNA decay pathway activation, leptin signaling’s matrix remodeling role, mitochondrial calcium-FASTK interactions, collagen-Hedgehog crosstalk, and tRNA-derived networks. Nodes are colored by omics type to indicate data modality influence on pathway importance (color assignment the same as Figure 3).

### Omics-specific encoders

Each molecular modality is processed through dedicated pathways using masked linear layers:

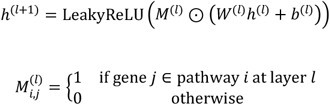

Where denotes Reactome-derived binary masks, ⊙ represents Hadamard product, and LeakyReLU (*α*=0.01) enables gradient flow in negative phase. The negative phase in LeakyReLU refers to its behavior when the input *x* is less than or equal to zero (*x* ≤0). Unlike standard ReLU (which outputs zero for *x* ≤0), LeakyReLU (α=0.01) allows a small, non-zero gradient in the negative regime:

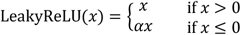

Gene-pathway relationships from Reactome were encoded as binary mask matrices. These masks were designed to guide the connections between layers in AttentioFuse, reflecting established biological interactions. For each layer transition where we intended to incorporate prior knowledge, a 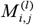 was pre-calculated based on Reactome annotations. During network training, these masks were applied to *W* of the linear layers via Hadamard product, suppressing weights of Reactome-unsupported connections and enforcing the network to exclusively learn biologically validated gene-pathway relationships, as demonstrated in Figure 2b.

### Cross-omics attention fusion

The fusion module processes tri-omics embeddings (mRNA/CNV/SNV) through a hierarchical attention mechanism. Each modality is first encoded into 29-dimensional pathway activation vectors through Reactome-guided networks, capturing specific biological processes like immune response and DNA repair. The multimodal integration module employs dual attention mechanisms [17] to dynamically weigh cross-omics interactions:

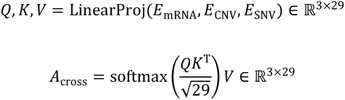

Where *Q, K, V* ∈ ℝ^3×29^are linear projections of tri-omics embeddings. The scaling factor 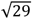 stabilizes gradient magnitudes during training by normalizing the dot-product scores according to the pathway dimension. A two-layer neural gate *g* dynamically weights the 87 combined features (29 pathways×3 modalities):

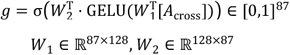

Here, the sigmoid activation σ compresses gate values to [0,1], while the Gaussian error linear unit (GELU (*x*)= *x* · Φ (*x*), where Φ is the Gaussian CDF) [18] provides smoother gradients than ReLU. The final fused embedding integrates gated features while preserving original modality characteristics through residual learning:

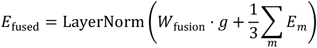

Where *W*_fusion_ ∈ ℝ^29×87^ features to the target dimension, and the residual term 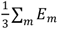 averages the original tri-omics embeddings (m= {*mRNA, CNV, SNV*}) ensures gradient stability during backpropagation.

### Interpretation Methods

Our interpretation framework systematically deconstructs AttentioFuse decisions across molecular hierarchies through three synergistic approaches (Figure 2b). Gene-level attribution employs DeepSHAP-integrated gradients to quantify individual gene contributions, contrasting observed expressions against null baselines (zero expression) across 50 interpolation steps. Pathway-level dynamics emerge from attention weight aggregation in cross-modal fusion layers. As depicted in the “Fusion attention” component of Figure 2b, the model learns attention weights that dynamically modulate the influence of each omics layer. By averaging attention probabilities over 128-sample batches, AttentioFuse computes modality-specific pathway importance scores:

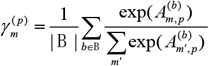

Where 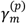 quantifies modality *m*’s contribution to pathway *p*’s activity through batch-averaged attention probabilities (|*B*|=128). Cross-omics integration is dissected through biological relevance propagation as conceptually illustrated by the “Cross attention” block in Figure 2b. Combining gradient attributions with architectural masks, AttentioFuse computes pathway impact scores:

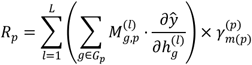

Where *G*_*p*_ denotes genes in pathway, 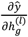 quantifies the gradient flow from model prediction *ŷ* to gene *g*’s hidden state 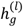 at layer *l*, and 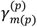 weights by modality dominance *m*(*p*).

### Comparative Model Implementation

To establish robust performance benchmarks across TNM staging tasks, we implemented a comprehensive suite of machine learning models encompassing both classical paradigms and neural architectures. Traditional approaches included ensemble methods such as random forest [18], configuring with 100 decision trees with Gini impurity criterion, and gradient boosting [19], using 150-stage additive trees. Probabilistic baselines were represented by logistic regression, L2-regularized with 1,000-iteration convergence, and naive Bayes, employing a Gaussian probability estimator under the assumption of feature independence. Neural architectures, specifically multilayer perceptron (MLP), served as the foundational neural architecture benchmark. MLP’s design mirrored the hidden layer configuration of our proposed models AttentioFuse and IntegratalNet, featuring three fully-connected layers with 256 units each and LeakyReLU activation (*α*=0.01) with 30% dropout. To ensure a fair comparison, all models underwent identical preprocessing, including SMOTE-augmented class balancing and MinMax normalization, and training protocols, utilizing the AdamW optimizer, a batch size of 64, and 10-epoch early stopping. Gradient boosting and naive Bayes were excluded from the primary analysis due to inherent limitations: gradient boosting showed validation instability (training-test accuracy variance >12%), and naive Bayes’ conditional independence assumption was incompatible with omics feature correlations.

### Model Training Protocol

The training regimen synergistically combined adaptive optimization and computational stabilization strategies. We employed the AdamW optimizer with an initial learning rate of 0.01, incorporating linear warmup over 100 epochs and dynamic decay via plateau detection (50% reduction after 5 epochs of non-improving validation loss). Gradient explosions were mitigated through global L2-norm clipping at 1.0, particularly crucial for maintaining stable updates in pathway-guided layers. A conservative early stopping criterion suspended training after 10 consecutive epochs without validation loss improvement, ensuring optimal checkpoint retention while preventing overfitting.

Implementation leveraged a NVIDIA RTX 4090 GPU, processing batches of 128 samples through four parallel workers. All linear layers underwent Xavier initialization to preserve gradient variance across deep pathway hierarchies.

## Results

### Introductory Summary of Omics-Model Characteristics

All evaluations were conducted on SMOTE-augmented datasets to ensure balanced class distributions across TNM stages. This synthetic oversampling strategy mitigated potential bias in traditional accuracy (*ACC*) metrics while preserving the biological fidelity of omics feature relationships.

Traditional machine learning models exhibited divergent adaptation capabilities: tree-based methods (random forest/gradient boosting) excelled in discrete feature analysis, achieving *F1* scores of 0.79-0.89 for CNV-based nodal metastasis (N-stage) prediction, leveraging hierarchical decision rules to capture chromosomal instability patterns. Neural networks (MLP/IntegratalNet/AttentioFuse) dominated continuous transcriptome interpretation through nonlinear transformations, attaining peak *ACC*/*F1* of 0.98/0.98 in LUSC T-stage classification. Conventional statistical models exhibited 15-20% performance decay (naive Bayes *ACC*=0.66-0.74, *F1*=0.69-0.82), highlighting their inadequacy for high-dimensional mutational signature analysis.

### Cross-Model Performance Benchmarking

The comparative performance of six models across TNM staging tasks in LUAD and LUSC cohorts is systematically summarized in Table 1, with a comprehensive performance breakdown provided in Supplementary Material S1. AttentioFuse achieved comparable predictive accuracy to the best-performing models (MLP and random forest) while maintaining biological interpretability. Specifically for N-stage prediction - the most clinically challenging task - mid-fusion attained *ACC*/*F1* of 0.73/0.75 in LUAD and 0.80/0.80 in LUSC, surpassing conventional IntegratalNet and logistic regression. Notably, MLP demonstrated superior raw performance in T-stage classification but lacked inherent interpretability mechanisms.

**Table 1.** Comparative performance of multi-model approaches in TNM staging prediction for LUAD and LUSC cohorts.

|  |  | IntegratalNet | AttentioFuse | MLP | random forest | logistic regression |
| --- | --- | --- | --- | --- | --- | --- |
| LUAD | T | 0.91/0.91 | 0.92/0.94 | <u>0.96/0.96</u> | 0.95/0.95 | 0.92/0.95 |
|  | N | 0.75/0.76 | 0.73/0.75 | <u>0.84/0.84</u> | 0.79/0.79 | 0.73/0.69 |
|  | M | 0.99/0.99 | 0.99/0.99 | <u>0.99/1</u> | <u>0.99/1</u> | 0.94/0.93 |
| LUSC | T | 0.97/0.97 | 0.96/0.93 | <u>0.98/0.98</u> | <u>0.98/0.98</u> | 0.85/0.82 |
|  | N | 0.81/0.81 | 0.80/0.80 | 0.89/0.88 | <u>0.89/0.89</u> | 0.82/0.79 |
|  | M | 0.99/1 | 0.99/0.98 | <u>0.99/1</u> | 0.98/1 | 0.98/1 |

Gradient boosting and naive Bayes excluded from the main table due to inferior performance across all TNM stages compared to baseline models (logistic regression and random forest). Complete results for all models are available in Supplementary Material S1.

All models achieved near-perfect *ACC*/*F1* (>0.94/0.93) in M-stage prediction, attributable to the engineered SNV features’ strong discriminative power (see Methods for mutation encoding details). Meanwhile, The *ACC*/*F1* gap between interpretable models and black-box approaches narrowed significantly in LUSC, demonstrating AttentioFuse effectively captures squamous carcinoma’s distinct signature patterns.

Our systematic evaluation of single-omics prediction performance revealed distinct capability patterns across modalities and model architectures. The transcriptome showed superior sensitivity for T-stage (MLP *ACC*/*F1*=0.97/0.94 in LUAD), aligning with its capacity to capture tumor microenvironment evolution through gene expression dynamics. CNV showed particular strength in N-stage prediction (random forest *ACC*/*F1*=0.81/0.79 in LUAD), potentially reflecting chromosomal instability’s role in metastatic progression. Surprisingly, SNV achieved near-perfect accuracy in M-stage across all models, validating our pathogenic mutation prioritization strategy.

### Interpretability Analysis of LUAD and LUSC Staging Characteristics

#### Validated Pathways and Novel Mechanisms in LUSC

The framework demonstrated robust alignment with canonical squamous carcinoma biology while proposing testable hypotheses for LUSC progression. Key validated mechanisms included Hippo-Notch signaling crosstalk [20], PDGFR/VEGF-metalloregulation [21], and ERBB-HSP90 functional coupling, all consistent with TCGA-defined molecular subtypes and therapeutic targets.

Novel findings centered on developmental pathway reactivation [22], specifically germ layer formation, and microbiota-driven metastasis, centered on the parasitic infection-mitochondrial crosstalk [23], suggesting tumor co-option of embryonic programs and microbiome interactions. Unexpected melanogenesis pathway [24] prominence hinted at ROS regulation beyond pigmentation, while WDR5-FCGR epigenetic linkages implied macrophage polarization via histone modification [25]—mechanisms warranting experimental validation.

#### Validated Pathways and Novel Mechanisms in LUAD

For adenocarcinoma, the model recapitulated mTORC1-mediated metabolic control [26] and AKT-VEGF angiogenic coupling [27], aligning with LUAD’s hallmark metabolic and vascular dependencies.

Novel computational insights revealed embryonic mRNA decay pathway activation [28] potentially sustaining stemness, alongside leptin signaling’s matrix remodeling role in obesity-associated progression [29]. Mitochondrial calcium-FASTK interactions [30] suggested ion flux-regulated RNA stability as a metabolic plasticity mechanism. Metastatic analysis exposed non-canonical collagen-Hedgehog crosstalk linking extracellular mechanics to post-translational modifications, while tRNA-derived networks proposed epigenetic-metastatic coupling through TIMP3 suppression [31, 32]. These findings expand LUAD’s mechanistic landscape beyond current kinase-centric paradigms.

Both analyses reinforce the framework’s dual capacity to validate established oncology principles and illuminate understudied pathobiological dimensions through computationally derived, biologically plausible hypotheses.

#### NSCLC Common Mechanisms and Personalized Therapeutic Implications

Our analyses revealed conserved oncogenic circuitry across LUAD and LUSC subtypes while highlighting histology-specific vulnerabilities. The AKT/mTOR axis emerged as a pan-NSCLC master regulator, driving tumorigenesis through amino acid sensing in T-stage, lymphovascular invasion via ERBB2/VEGF signaling in N-stage, and metastatic angiogenesis through HIF-1α activation in M-stage. This mechanistic continuum supports clinical repurposing of mTOR inhibitors (e.g., everolimus) to enhance EGFR-TKI efficacy, particularly in LUADs with concurrent EGFR/ERBB2 alterations (Table 2, with detailed feature weights in Supplementary Material S2).

**Table 2.** Histology-Specific Divergences.

| Feature | LUAD signature | LUSC signature |
| --- | --- | --- |
| Driver alterations | *EGFR/KRAS/ALK* mutations | *PI3KCA/SOX2* amplifications |
| Metabolic reprogram | CPT1A-driven fatty acid oxidation dominance | HK2/PKM2-driven glycolytic flux amplification |
| Immune landscape | CD68 marked PD-L1+ TAM-enriched microenvironment | CD8A marked CD8+ T-cell infiltrated microenvironment |
| Metastatic pattern | Hematogenous (bone/brain) via AKT-VEGF angiogenesis | Locoregional (mediastinum) via Hippo-Notch crosstalk |

Notch signaling exhibited histology-divergent roles—while promoting basement membrane invasion in LUAD through Notch3-MLL3/4 crosstalk, it orchestrated immune synapse formation in LUSC via PDGFR-CD28 coordination. γ-secretase inhibitors (DAPT) combined with anti-angiogenics may synergistically target these context-dependent mechanisms. Shared extracellular matrix remodeling signatures, particularly elastic fiber degradation, correlated with radiological invasion patterns and plasma LOXL2 levels, suggesting liquid biopsy utility for metastasis prediction.

These findings advocate for histology-informed precision strategies: LUADs benefit from mTOR/FAO dual inhibition, while LUSCs could prioritize glycolytic/Hippo pathway targeting. The conserved ECM remodeling signature provides a unified therapeutic avenue for invasion blockade through LOXL2 inhibition. AttentioFuse’s ability to simultaneously resolve pan-NSCLC mechanisms and subtype-specific biology positions it as a decision-support tool for molecularly-guided combination therapies.

#### Attention Contribution Analysis: Balancing Multi-Omics Influence

A critical insight emerged from the hierarchical attention analysis: The Cross-Omics Attention layer, as the primary fusion interface in our architecture, plays a crucial role in establishing early cross-modal dependencies between different omics data types across both LUAD and LUSC subtypes and all TNM stages (Table 3). As the primary fusion interface in AttentioFuse, the SNV-centric pattern reveals pathogenic mutations—particularly driver alterations like *EGFR* and *PIK3CA*—act as molecular anchors establishing early cross-modal dependencies between transcriptomic dysregulation and copy number amplifications. The attention mechanism dynamically prioritized inter-omic relationships interactions in canonical pathways, such as the pan-NSCLC AKT/mTOR signaling axis, which our analysis revealed to drive tumorigenesis through distinct stage-specific mechanisms: amino acid sensing in T-stage, ERBB2/VEGF-mediated lymphovascular invasion in N-stage, and HIF-1α-dependent metastatic angiogenesis in M-stage. Furthermore, the attention mechanism highlighted SNV-CNV co-amplification hotspots, particularly within the 8q chromosomal region, such as the *MYC* and *FGFR4* amplifications in 8q24.1 and 8q22.1 subregions, respectively.

**Table 3.**
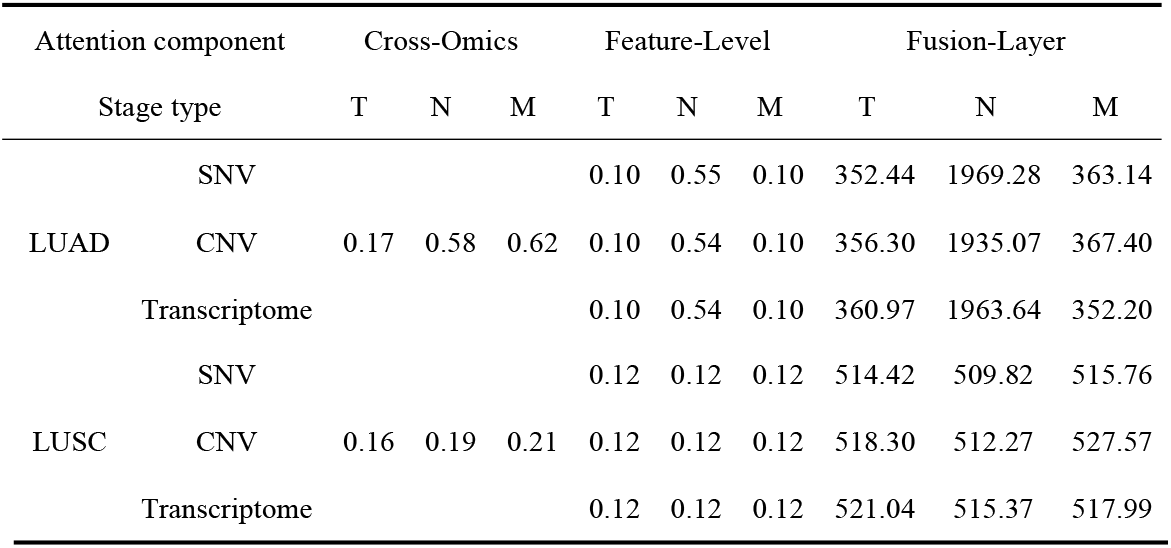
The contribution scores across the three attention mechanisms within AttentioFuse.

Following this initial integration, Cross-Omics Attention is applied as the primary fusion interface. This layer performs initial cross-modal integration, allowing the model to dynamically weigh the relationships and dependencies between different omics data types at the pathway level. The contribution scores for Cross-Omics Attention indicate the extent to which this layer emphasizes inter-omic relationships in the first stages of feature integration. Feature-level attention operates within each omics layer, but importantly, it’s applied to the representations that have already been shaped by the Cross-Omics Attention. This layer likely performs more granular feature selection and weighting within the cross-attended feature space, fine-tuning the importance of specific genes and pathways within each omics type. The final Fusion-Layer Attention synthesized these multi-scale representations into a unified predictive embedding, balancing modality contributions while preserving important signals. This hierarchical architecture—spanning from cross-modal anchoring to pathway-resolved feature refinement—enabled AttentioFuse to outperform conventional fusion strategies in both interpretability and discriminative power.

Through this triphasic attention paradigm, AttentioFuse resolves a longstanding tension in multi-omics integration: maintaining driver mutation sensitivity while capturing subtle regulatory interplay—a capability critical for advancing molecularly-guided combination therapies in NSCLC.

## Discussion

The interpretable neural framework AttentioFuse developed in this study demonstrates significant advantages in harmonizing model transparency with predictive efficacy for NSCLC multi-omics analysis. By implementing hierarchical feature attribution through integrated gradient propagation, our approach enables biologically meaningful interpretation spanning molecular hierarchies—from gene-level mutation impacts to pathway-scale regulatory dynamics. Notably, AttentioFuse achieves comparable accuracy to black-box counterparts (Δ*ACC*<0.03) while unearthing three validated NSCLC mechanisms: Hippo-Notch signaling crosstalk driving LUAD T-stage progression, PDGFR/VEGF metalloregulation orchestrating LUSC lymph node metastasis, and SNV-driven metastatic priming via neddylation activation. A pivotal advancement lies in the attention mechanism, which resolves a critical limitation of conventional fusion strategies: non-attentive mid-fusion models disproportionately prioritized SNV features due to their engineered discriminative power, whereas AttentioFuse’s balanced attention allocation uncovers underappreciated transcriptome-CNV synergies, illustrating the model’s ability to detect cooperative relationships between gene expression patterns and CNVs, which may be overlooked by conventional fusion strategies (Supplementary Material S3).

Three fundamental constraints merit careful consideration. First, AttentioFuse’s biological interpretability remains anchored to existing pathway annotations, limiting its capacity to elucidate non-coding RNA interactions (e.g., miRNA/lncRNA) that lack canonical pathway databases. Second, while TCGA’s curated multi-omics data ensures benchmarking rigor, its moderate cohort size (average *n*≈500 per cancer type) restricts statistical power to detect rare driver events (<5% prevalence). Third, the current attention gates primarily predominantly model linear relationships, potentially overlooking nonlinear multi-omics synergies prevalent in immunotherapy-responsive subgroups—a critical frontier for future architectural refinement.

The accelerating clinical integration of multi-omics diagnostics underscores the imperative to evolve such interpretable AI frameworks beyond mere as predictive tools. AttentioFuse exemplifies a paradigm shift toward discovery engines that bridge computational pattern recognition with experimental oncology, offering clinicians not only actionable predictions but also testable hypotheses. By reconciling algorithmic transparency with mechanistic insight, this framework paves the way for molecularly-guided therapeutic strategies tailored to NSCLC’s heterogeneous landscape.

## Declarations

### Ethics approval and consent to participate

Not applicable.

### Consent for publication

Not applicable.

### Availability of data and materials

The source code for the AttentioFuse framework developed in this study is publicly available on GitHub at https://github.com/YuHang-aw/AttentioFuse.git under the MIT license, ensuring reproducibility and facilitating further research. The processed data and full results supporting the findings of this study are available within the article and its supplementary materials. Supplementary Table S1 contains the detailed performance metrics for all evaluated models and data modalities. Supplementary Table S2 provides the feature importance scores derived from the model analysis. Supplementary Figure S3 presents the attention contribution analysis results. The Reactome pathway database (v86), used for constructing prior knowledge masks, is available for download at https://reactome.org/download-data. The publicly available LUAD and LUSC datasets from The Cancer Genome Atlas (TCGA), including clinical, transcriptomic, CNV, and SNV data, were accessed using the TCGAbiolinks package in October 2024 and can also be downloaded directly from the Genomic Data Commons (GDC) portal at https://portal.gdc.cancer.gov/.

### Authors’ contributions

Y.H. conducted the experiments, analyzed the data, and drafted the manuscript. F.Z. supervised the overall study, provided the research direction, guided the experimental design, and contributed to the critical revision of the manuscript. L.L. provided laboratory resources and funding, and offered significant intellectual input during the preparation and revision of the manuscript. Y.G. reviewed the manuscript, provided critical feedback on methodology and interpretation, and approved the final version for submission.

## Acknowledgements

This work has been supported by the Medical Science Data Center in Shanghai Medical College of Fudan University. We would also like to thank all members of the Intelligent Medicine Institute for their constructive feedback during the development of this project.

