## Supplemental Data 3 for "From Black Box to Biological Insight: AttentioFuse Unlocks Multi-Omics Dynamics in Lung Cancer"

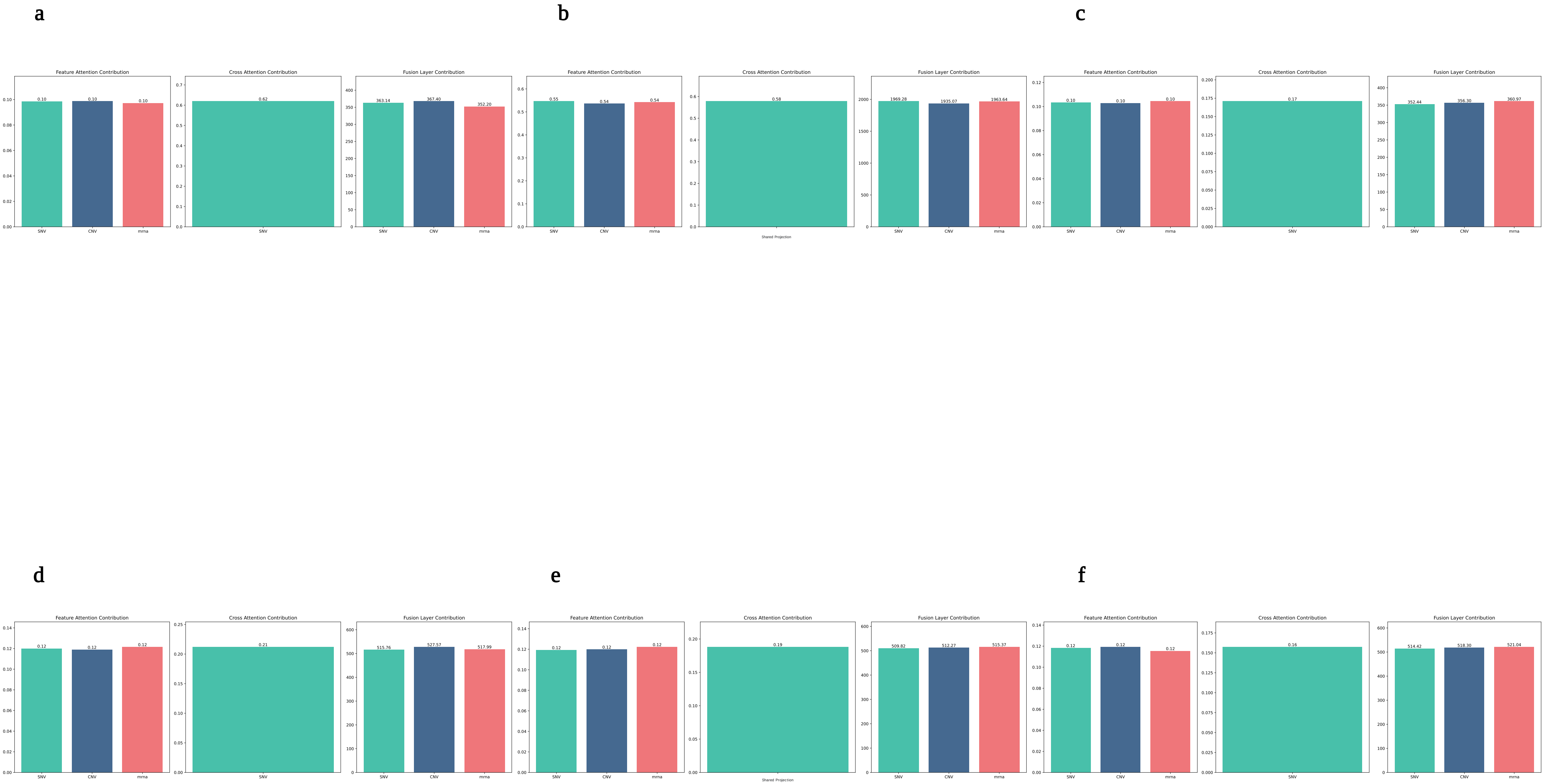


Supplementary Figure S3. AttentioFuse Model – Deconstructing Attention Contributions. This figure visualizes the contribution of the three attention mechanisms within the AttentioFuse framework for each omics data type and TNM staging task. Panels (a)-(c) show the "Feature Attention Contribution," "Cross Attention Contribution," and "Fusion Layer Contribution" for Lung Adenocarcinoma (LUAD) in (a) M-stage, (b) N-stage, and (c) T-stage prediction. Panels (d)-(f) present the same attention analysis for Lung Squamous Cell Carcinoma (LUSC) for (d) M-stage, (e) N-stage, and (f) T-stage. Within each panel group, the "Sum of Absolute Weights" (Y-axis) is shown for SNV (green), CNV (blue), and Transcriptome (red, labeled as 'mrna' - shorthand for Transcriptome). The three attention types represent: Feature-Level Attention (within-omics feature importance), Cross-Omics Attention (initial inter-omic interaction); note that while the plots may visually emphasize SNV due to plotting order, the Cross-Omics Attention mechanism uses *shared weights* and involves *all three* omics types), and Fusion-Layer Attention (overall omics layer weighting at fusion).
